# The Link to Oxidative Metabolism Varies across rs-fMRI Metrics: A Whole-Brain Assessment Using Macrovascular Correction

**DOI:** 10.1101/2025.02.11.633958

**Authors:** Xiaole Z. Zhong, Hannah Van Lankveld, J. Jean Chen

## Abstract

One of the major obstacles to the clinical application of resting-state functional magnetic resonance imaging (rs-fMRI) is the complex nature of its measurements, which limits interpretability. An approach to enhance the interpretability of the rs-fMRI metrics is to link them to more fundamental brain physiology, especially cerebral metabolism. Previous studies have established associations between glucose metabolism (CMR_glu_) and rs-fMRI measurements. In spite of this, oxidative metabolism (CMRO_2_) is more closely related to cerebral blood flow (CBF) and thus the BOLD signal, and its relationship with CMR_glu_ is complex. Additionally, most currently published rs-fMRI metrics are uncorrected for macrovascular contribution, which may obscure the neuronal contributions. In this study, we measured resting CMRO_2_ (along with the oxygen extraction fraction, OEF and cerebral blood flow, CBF) using gas-free calibrated fMRI. We used linear mixed-effects (LME) models to examine associations between CMRO_2_ and various rs-fMRI metrics before and after macrovascular correction. We found that: 1) significant associations exist between CMRO_2_ and multiple rs-fMRI metrics, with the strongest association found for the global functional density (gFCD) and the weakest for seed-based functional connectivity (FC); 2) associations with rs-fMRI metrics also varied for OEF and CBF; 3) significant sex differences were observed in the above associations; 4) the use of macrovascular correction substantially strengthened the goodness fit of all LME models examined. This latter improvement further validates the use of macrovascular correction in rs-fMRI. These results provide a framework for linking rs-fMRI metrics to fundamental brain physiology, thus improving interpretability of rs-fMRI measurements. This is the first study to formally link whole-brain MRI-based baseline CMRO_2_ and rs-fMRI metrics, and helps to push the envelope for rs-fMRI in future clinical applications.

## Introduction

Resting-state functional magnetic resonance imaging (rs-fMRI), most commonly based on the blood-oxygenation level-dependent (BOLD) contrast, aims to use spontaneous brain activity (B. Biswal et al., 1995) to investigate brain function in health and disease. Most rs-fMRI studies lend themselves to connectivity analysis, originally based on Pearson’s correlation (B. Biswal et al., 1995) and later developed into a family of connectivity metrics. Moreover, the resting-state fluctuation amplitude (RSFA) has been used as a means of understanding the temporal variability of rs-fMR (Kannurpatti et al., 2011). Recently, non-linear metrics, notably entropy, have revealed novel insight into the relationship between the rs-fMRI signal complexity and cognitive performance (Wang, 2021). In spite of the numerous metrics that have been proposed as markers of neuronal function, there has been limited application in clinical practice, due in part to the lack of understanding of the neuronal mechanism that underlies the rs-fMRI signal (Lee et al., 2013). For instance, many of these metrics have significant vascular contributions that may obscure the metabolic interpretations (vice versa).

A natural approach to investigating this question of neuronal mechanisms of rs-fMRI is to relate to task-based fMRI. Local neuronal specificity has been investigated by linking fMRI to its neurovascular substrates, including cerebral blood flow (CBF) and cerebral metabolic rate of oxygen (CMRO_2_), which are believed to be more directly associated with local neural activity (Davis et al., 1998). Typically, a reduction in OEF caused by an increase in CBF in response to a CMRO_2_ increase is associated with an increase in BOLD fMRI during local neural activation. Such associations are also relevant to understanding rs-fMRI if the same dynamic coupling for tasks is assumed to apply at rest. This has led to previous research suggesting that functional networks can be derived not only from fluctuations in rs-fMRI but also from CMRO_2_ and CBF (B. B. Biswal et al., 1997; Wu et al., 2009). However, this approach assumes a dynamic neurovascular coupling similar to a steady-state coupling, which has not been established. Treating resting-state as a form of steady-state, an association between RSFA and baseline CBF by simulations and in-vivo measurements has been reported (Chu et al., 2018). Nevertheless, the relationship between rs-fMRI connectivity and steady-state resting CMRO_2_ has not yet been fully established. While this relationship is understandably distinct from that in task or dynamic state, it is nevertheless an important step in understanding and interpreting rs-fMRI metrics.

A seminal study by Raichle and colleagues using positron-emission tomography (PET) demonstrated that baseline CMRO_2_ is distributed heterogeneously across the resting brain, being particularly high in a set of functionally connected regions constituting the default mode network (DMN) (Raichle et al., 2001). In light of the high metabolism of the DMN, it is evident that this region is the most active in the resting state. This evidence encourages the investigation of neurometabolism in other resting-state brain networks. Although such investigations have been limited, the cerebral metabolic rate of glucose (CMR_glu_) has also been used to establish links between rs-fMRI and brain metabolism at rest. As an example, a positive association has been established between CMR_glu_ and functional connectivity density (FCD) across different brain regions (Shokri-Kojori et al., 2019; Tomasi et al., 2013), where hubs regions, which were highly connected locally and globally, were associated with higher CMR_glu_ (Tomasi & Volkow, 2010).

CMR_glu_, however, is not directly related to the rs-fMRI signal, which is based on the BOLD effect and thus more directly related to CMRO_2_ (Davis et al., 1998; Hoge et al., 1999; Ogawa et al., 1993). Despite the fact that the majority of glucose undergoes complete oxidative phosphorylation, the metabolism of glucose and oxygen are not always coupled (Gibbs et al., 1942). Moreover, aerobic glycolysis, defined as a failure to fully metabolize glucose despite having sufficient oxygen (under-consumption of oxygen), is also found in the healthy brain, with the highest incidence located within the DMN (Vaishnavi et al., 2010). The metabolism of oxygen and glucose are decoupled in both cases described above, which brings us back to the importance of the initial study of CMRO_2_ by Raichle (Raichle et al., 2001), which, however, did not explicitly involve rs-fMRI metrics.

Establishing a connection between rs-fMRI and metabolism would assume that rs-fMRI metrics are dominated by neuronal sources. However, the BOLD signal can be significantly affected by physiological factors that are not neuronal, which makes it more difficult to quantify the rs-fMRI metrics. The typical frequency range of neural activity is 0.01 to 0.1 Hz (Tong, Hocke, et al., 2019), though time-lag spatiotemporal correlational structures are also observed among low-frequency oscillations (sLFOs) within this frequency range (Tong et al., 2015). It is also found that the sLFOs are especially prominent in the region around the macrovasculature, especially the large vein (Zhong, Tong, et al., 2024), contributing to robust correlation patterns (Tong, Yao, et al., 2019). During the early days of fMRI, macrovasculature (defined as major arteries and sinuses) was found to have a greater influence than microvasculature (defined as capillaries, arterioles, and venules) (Boxerman et al., 1995). The same is true for higher fields (Menon, 2012), even as the sensitivity to microvasculature increases (Menon & Goodyear, 1999). In fact, rs-fMRI has been demonstrated in detecting venous structure in previous studies on rats (Hyde & Li, 2014), and at 11.7 T, the venous BOLD response is twice as high as that in brain tissue (Yu et al., 2012). In our recent study (Zhong, Tong, et al., 2024), we found that the macrovasculature consistently exhibited high correlations within itself and that these correlations are responsible for a considerable portion of gray-matter-macrovasculature correlations. Moreover, the contribution extends well beyond the boundaries of macrovasculature, with a substantial contribution found at up to 1.2 cm from macrovasculature boundaries, which increases the difficulty of correcting macrovascular bias. Thus, the macrovascular confound is non-trivial, and its effect on the neuronal interpretation of rs-fMRI can be profound but is currently unknown. Recently, we proposed a novel method to correct for such macrovascular effects using biophysical modelling (Zhong, Polimeni, et al., 2024), but its effect on the interpretability of rs-fMRI metrics has yet to be demonstrated. We think that baseline physiological metrics such as CMRO_2_ can help establish such an interpretation, as described earlier.

In this study, we hypothesize that different rs-fMRI metrics are indeed associated with CMRO2 but to different extents, and that such associations can be enhanced by our novel macrovascular correction. A better understanding of associations between macrovascular-corrected rs-fMRI metrics and fundamental physiological variables could provide a solid basis for interpretation of rs-fMRI findings.

## Methods

### Theory: gas-free calibrated fMRI

In this study, the metabolic metrics OEF and CMRO_2_ were calculated using gas-free calibrated fMRI, which is based on the Yablonskiy random-cylinder model (Hirsch et al., 2014; Yablonskiy & Haacke, 1994). The OEF was estimated based on,

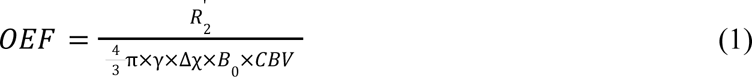

where *R*_2_’ is estimated from *R*_2_ and *R*_2_* as will be described in the subsection titled *Baseline metabolism estimation*, and reflects MR signal dephasing resulting from local magnetic field inhomogeneities. γ is the proton gyromagnetic ratio, and Δχ is the susceptibility difference between oxygenated and deoxygenated blood, CBV is cerebral blood volume, and its estimation will be described in the *Baseline perfusion estimation* section. OEF was restricted to be between 0 and 1. Using the OEF estimate, quantitative CMRO_2_ can then be estimated (Göttler et al., 2019) as follows,

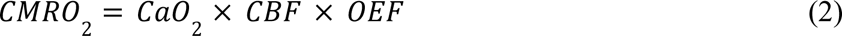

where CaO_2_ is the arterial oxygen content, set to 19 *ml O* /100 *ml blood* (An et al., 2001).

### Participants

This study involves 20 young healthy participants (10 males and 10 females between the ages of 20 and 32 years). No participant reported having a history of cardiovascular disease, psychiatric illness, neurological disorder, malignant disease, or medication use that may have affected the study. Participants were recruited through the Baycrest Participants Database, which consists of residents from Baycrest and surrounding communities. The study was approved by the research ethics board (REB) of Baycrest, and the experiments were performed with the understanding and written consent of each participant, according to REB guidelines.

### MRI acquisition

The images were acquired using a Siemens Prisma 3 Tesla System (Siemens, Erlangen, Germany), which employed 20-channel phased-array head coil reception and body coil transmission. The following data were acquired for each participant: (i) T1-weighted structural image (sagittal, 234 slices, 0.7 mm isotropic resolution, TE = 2.9 ms, TR = 2240 ms, TI = 1130 ms, flip angle = 10°); (ii) two time-of-flight (TOF) scans, with coronal and sagittal flow encoding, respectively (coronal: 0.8 x 0.8 x 2.125 mm thickness, 100 slices, TR = 16.6 ms, TE = 5.1 ms, flip angle (α) = 60°; sagittal: 0.8 x 0.8 x 2.125 mm thickness, 80 slices, TR = 16.6 ms, TE = 5.1 ms; flip angle (α) = 60°); (iii) one dual-echo (DE) pseudo-continuous arterial spin labelling (pCASL) (courtesy of Danny J. J. Wang, University of Southern California) for recording CBF and BOLD dynamics (TR = 4.5 s, TE1 = 9.8 ms, TE1 = 30 ms, post-labelling delay = 1.5 s, labelling duration = 1.5 s, flip angle (α) = 90°, 3.5 mm isotropic resolution, 35 slices, slices gap = 25%, scanning time = 4 minutes), complete with a M_0_ scan in which the TR was 10 s and the scan time 45 s, all other parameters remaining the same; (iv) one multi-echo gradient echo (MEGRE) scan for R_2_ mapping (TR = 4 s, TE ranged from 5 ms to 60 ms with the step of 5 ms, flip angle = 20°, magnitude and phase recording, same spatial resolution as the pCASL); (v) SE MRI data for R_2_ mapping (TR = 10 s, TE =[17, 33, 50, 66, 91, 107, 116] ms, same spatial resolution as the pCASL). The participants were imaged while viewing naturalistic stimuli to minimize random mind-wandering, thereby enhancing the reproducibility of the results (Gal et al., 2022).

### Macrovascular correction

An overview of the BOLD signal simulation framework can be found in our previous study (Zhong, Polimeni, et al., 2024). Based on our previous comparison, the macro-VAN model was selected for this simulation to provide a more accurate estimation of macrovascular BOLD signals, particularly from venous vessels.

### Blood-vessel segmentation

The first step for macrovascular correction is the creation of a vascular mask. The strategies for segmenting and processing macrovasculature are summarized in Figure 1a. For each subject, TOF data were registered to T1 space (FSL MCFLIRT, DOF = 6, cost function = correlation) and then segmented using the BrainCharter Toolbox (Bernier et al., 2018). Visual inspection was conducted to ensure that the macrovasculature segmentations were free of artifacts (e.g., wrap-around artifacts). The vascular segmentations from both encoding directions were summed and then binarized to produce the final segmentation. The final segmentations were first upsampled to a 0.175 mm (i.e. 0.7 mm/4) isotropic resolution (required for the simulation; see our previous work (Zhong, Polimeni, et al., 2024)) and then downsampled to the rs-fMRI resolution (3.5 mm resolution). The blood-volume fraction (fBV) was estimated by counting the number of high-resolution voxels (0.175 mm isotropic resolution) occupied by vessels in each rs-fMRI voxel (3.5 mm isotropic resolution with a 25% slice gap). As the TOF data contained arteries, veins and false-positive detections, they had to be manually identified and separated into separate maps using the anatomical atlas (Tortora & Derrickson, 2018). The resulting venous vasculature from a representative data set is illustrated in **Figure 2**. In this work, large veins are defined as vessels of sizes similar to that of the superior sagittal sinus, the straight sinus, the inferior sagittal sinus, and the transverse sinus.

**Figure 1.**
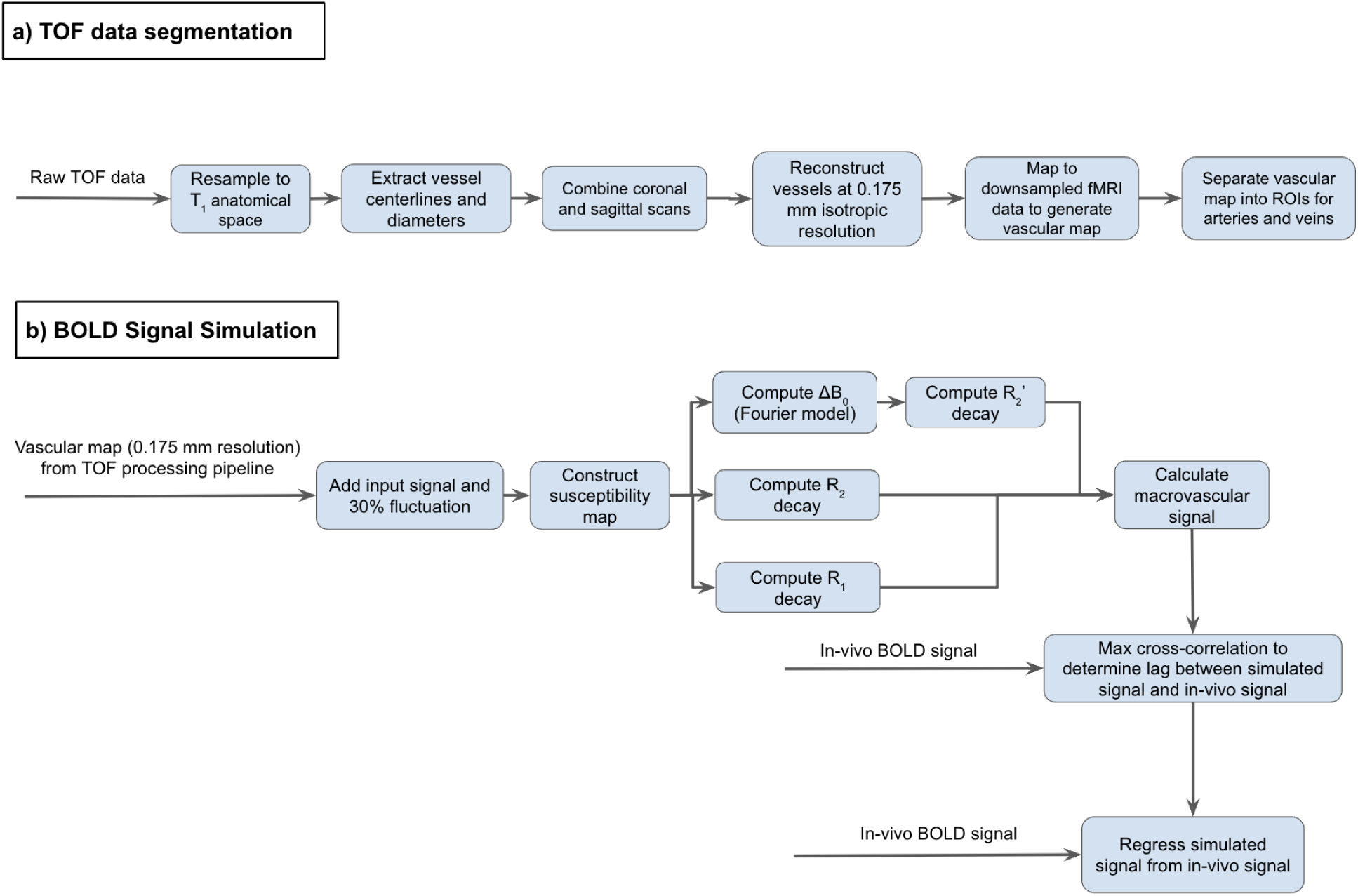
Overview of the macrovascular correction procedure. a) the TOF data preprocessing and segmentation pipeline; b) BOLD signal simulation and regression.

**Figure 2.**
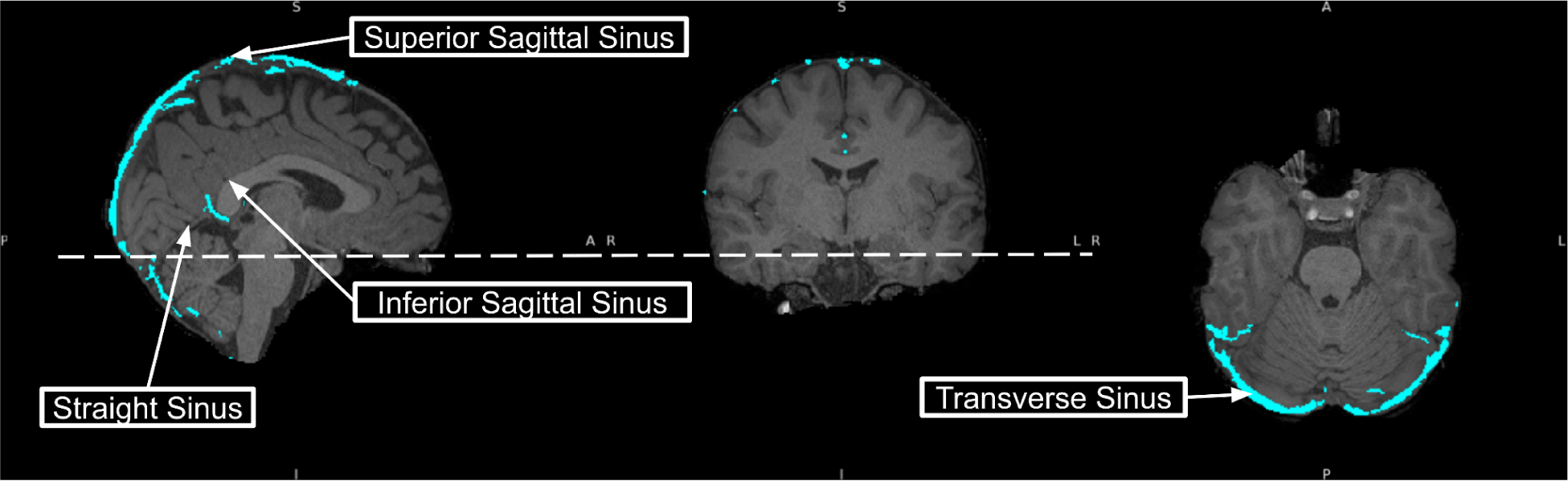
Venous vasculature demonstration overlays the anatomical images and defines macrovsculatue in this study (data shown from a representative data set). Blue is the vein segmented from the TOF data overlay on the T1 anatomical image; the white dashed line indicates the axial slice position.

### Macrovascular signal simulation and correction

A diagram summarizing the simulation steps is shown in **Figure 1b**. We used the same simulation process as in our previous study (Zhong, Polimeni, et al., 2024), adapted to the currently used spatial resolution with the exception that the input sinusoid was replaced with the in-vivo BOLD data. This was chosen to outperform the sinusoid in terms of accounting for signal variability in in-vivo BOLD, and also as it is less likely to include regional neuronal activity than the global signal (Tong, Yao, et al., 2019). The in-vivo BOLD input was generated as the square root of the in-vivo BOLD signal from the voxel with the maximum fBV as the input signal and then is demeaned and normalized by the maximum of the signal.

The *Supplementary Materials* contain additional information about the simulation procedure and mathematical model. Sinc*e* the simulated data would not incorporate any time lag among the in-vivo voxels, cross-correlation was applied between each in-vivo signal and simulated signal to determine the lag, leading to the maximum cross-correlation coefficient. This lag is considered the time difference between the simulated and in-vivo signals for specific voxels. The lag-corrected simulated signal was used as a nuisance regressor in a linear regression model on a voxelwise basis to remove the apparent macrovascular BOLD contribution. Signals thus corrected are referred to as the “post-correction” signal.

### rs-fMRI preprocessing and analysis

The rs-fMRI data was processed using two different pipelines: one for investigating seed-based functional connectivity (FC) and the other for seed-independent measures. Preprocessing steps shared by both pipelines include rejecting the first five volumes of BOLD data to allow the brain to enter steady state. Moreover, all metrics below were averaged within each functional network listed in the *Seed-based rs-fMRI analysis* subsection to calculate network-wise metrics.

### Seed-based rs-fMRI analysis

rs-fMRI connectivity analysis was performed using the CONN toolbox (Whitfield-Gabrieli & Nieto-Castanon, 2012) with the default_MNI preprocessed pipeline (bandpass filter: 0.01-0.1 Hz; regressed confounds: white matter, CSF and estimated subject-motion parameters). The networks of interest include the visual network (VN), sensorimotor network (SMN), the default mode network (DMN), the language network (LN), the salience network (SN), the dorsal attention network (DAN), and the frontoparietal network (FPN). To define each network, p-value maps for different sources were combined by selecting a minimum p-value and then thresholding by p < 0.0001. Network-wise seed-based FC was calculated by averaging the magnitude of correlation coefficients within each network.

### Seed-independent rs-fMRI analysis

rs-fMRI preprocessing pipeline was implemented with customized procedures based on tools from FSL (Jenkinson et al., 2012), AFNI (Cox, 1996) and FreeSurfer (Fischl, 2012). The following steps were included in the preprocessing steps: (a) 3D motion correction (FSL MCFLIRT), (b) slice-timing correction (FSL slicetimer), (c) brain extraction (FSL bet2 and FreeSurfer mri_watershed), (d) rigid body coregistration of functional data to the individual T1 image (FSL FLIRT), (e) regression of the mean signals from white-matter (WM) and cerebrospinal fluid (CSF) regions (fsl_glm), (f) bandpass filtering to obtain frequency band 0.01-0.1 Hz (AFNI 3dBandpass), and (g) spatial smoothing with a 6 mm full-width half-maximum (FWHM) Gaussian kernel (FSL fslmaths).

Once the BOLD signal had been preprocessed, we computed the following metrics:

1. Resting-state fluctuation amplitude (RSFA), calculated using the standard deviation of the BOLD signal normalized by the mean of the BOLD signal.
2. BOLD signal entropy, calculated using sample entropy with m = 3 and r = 0.6 related to the standard deviation as proposed by the previous study (Nezafati et al., 2020).
3. lFCD, calculated using the AFNI (3dLFCD) (Cox, 1996) with a threshold of 0.6, and the neighbourhood data was defined to include face-touching 6 voxels, as suggested by the previous study (Tomasi & Volkow, 2010).
4. gFCD, calculated using the same threshold value (0.6) but within the entire gray matter (Tomasi & Volkow, 2011).

Since density-related measurements follow an exponential distribution, log-transformed data was used for both lFCD and gFCD to transform data to a normal distribution (Shokri-Kojori et al., 2019).

### Baseline perfusion estimation

First, the ASL data was skull-stripped, and then the data was split into labelled and controlled images based on odd and even volumes. For both the controlled and labelled data, a motion correction was applied before surround subtraction was performed to determine the signal difference between the labelled and controlled images. To ensure the brain entered a steady state, the first five volumes of ASL data were also discarded, which is similar to the rs-fMRI process. The CBF value was quantified using the BASIL toolbox from FSL (oxford_asl) (Chappell et al., 2009). The CBF maps were also averaged within each functional network listed in the rs-*fMRI connectivity* subsection to calculate network-based metrics.

Fractional CBV will be calculated from CBF by applying linear regression between regionally-averaged baseline CBF (as ml/100ml/min) and CBV (as a fraction) values in gray and white matter (Eq. 3), measured by previous research with ^15^O steady-state inhalation PET in 34 healthy participants (Leenders et al., 1990; Zhang et al., 2018).

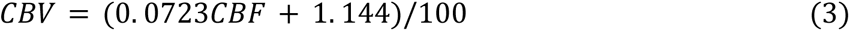

### Baseline metabolism estimation

Both the SE images and the MEGRE image were first skull-stripped, and then the rigid-body coregistration was performed to align SE and MEGRE images with the T1-weighed image. Using SE images, R_2_ (refocusable transverse relaxation rate) values were determined based on a single exponential model fitted to each TE. Since the MEGRE scan is sensitive to background fields, a background field correction was performed on the MEGRE data, which calculated the correction factor based on phase components of the MEGRE image and RF pulse shape to estimate gradients in the x, y, and z directions (Kaczmarz et al., 2020). The corrected MEGRE magnitude images were then fitted using a single exponential model to estimate R_2_ (non-refocusable transverse relaxation rate). The R_2_’ was calculated as the difference between R_2_ and R_2_ . The OEF and CMRO_2_ were estimated using R_2_’ and perfusion parameters as described in theory. As before, the OEF and CMRO_2_ values were also averaged within each functional network listed in the rs-*fMRI connectivity* subsection to calculate network-wise metrics.

### Statistical analysis

We used a linear mixed-effect model (LME) approach to assess the association between network-averaged CMRO_2_ and related physiological metrics and rs-fMRI metrics, both before and after macrovascular correction. In the LME, sex was also considered as a fixed effect of interest since significant sex effects were observed in both CBF (Mazzucco et al., 2024) and FC metrics (Tomasi & Volkow, 2012). The z-transform was applied to all parameters before fitting. For each fitting, outliers were detected and removed according to the 1.5 interquartile range (IQR) criterion. An overview of the parameters of interest is presented in **Table 1** and the model is described as **Eq. 4**. A significance level of 0.05 was used with a false discovery rate (FDR) correction for results of pre- and post-macrovascular correction separately. The r^2^ values are presented with the number of coefficients adjusted.

**Table 1.** Parameters for linear mixed effect model.

| $Y \sim 1 + X + \text{Sex} + X:\text{Sex}$ (Eq. 4) | |
| --- | --- |
| Y (rs-fMRI) | X (Metabolism and Hemodynamics) |
| RSFA | CMRO<sub>2</sub> |
| Seed-based FC | CBF |
| gFCD | OEF |
| IFCD |  |
| Entropy |  |

In order to better understand the inter-regional variability, a correlation analysis was performed between rs-fMRI metrics and BOLD physiological origins from seven networks for each participant with a significant threshold at p<0.05.

## Results

### Variations across networks

Network definitions varied depending on the rs-fMRI-based FC values, which were altered by macrovascular correction. This resulted in pre-correction and post-correction sets of network ROIs. **Figure 3** illustrates that there were no notable differences across networks in terms of CMRO_2_, nor are there for OEF and CBF. There were no significant differences in rs-fMRI metrics across all seven networks (the differences were smaller than the intra-network standard deviation) for either seed-based or seed-independent metrics, despite the appearance of heterogeneity in the values across networks.

**Figure 3.**
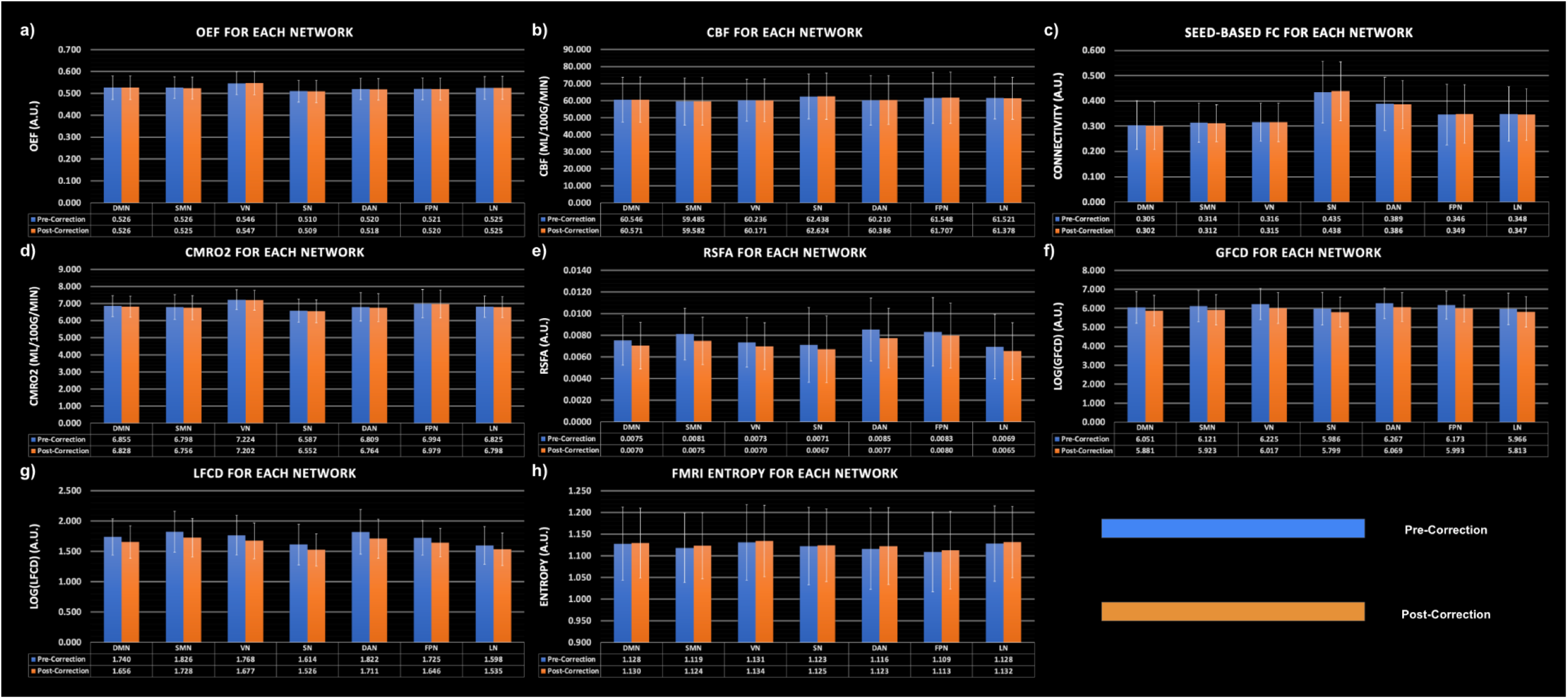
The average CMRO_2_, OEF, CBF and rs-fMRI metrics using network ROIs are defined based on rs-fMRI data pre- and post-macrovascular correction. The rs-fMRI metrics corresponding to the post-correction ROIs are of course corrected for macrovascular contributions. Grouped bars from left to right: DMN, SMN, VN, SN, DAN, FPN, and LN. Blue: pre-macrovascular correction and orange: post-macrovascular correction. Error bars represent standard deviation.

### Association between rs-fMRI metrics and physiology

We will first focus on the findings of post-macrovascular correction. The findings in this section are determined across all functional networks. As shown in **Figure 4a**, all seed-independent rs-fMRI metrics exhibited strong associations with CMRO_2_, however, no seed-based FC was associated with CMRO_2_. Segregating by the direction of the associations, gFCD, lFCD, and RSFA were positively associated with CMRO_2_, while rs-fMRI entropy was negatively associated with CMRO_2_. In comparison to the rest of the rs-fMRI metrics, the gFCD showed a stronger association with CMRO_2_.

**Figure 4.**
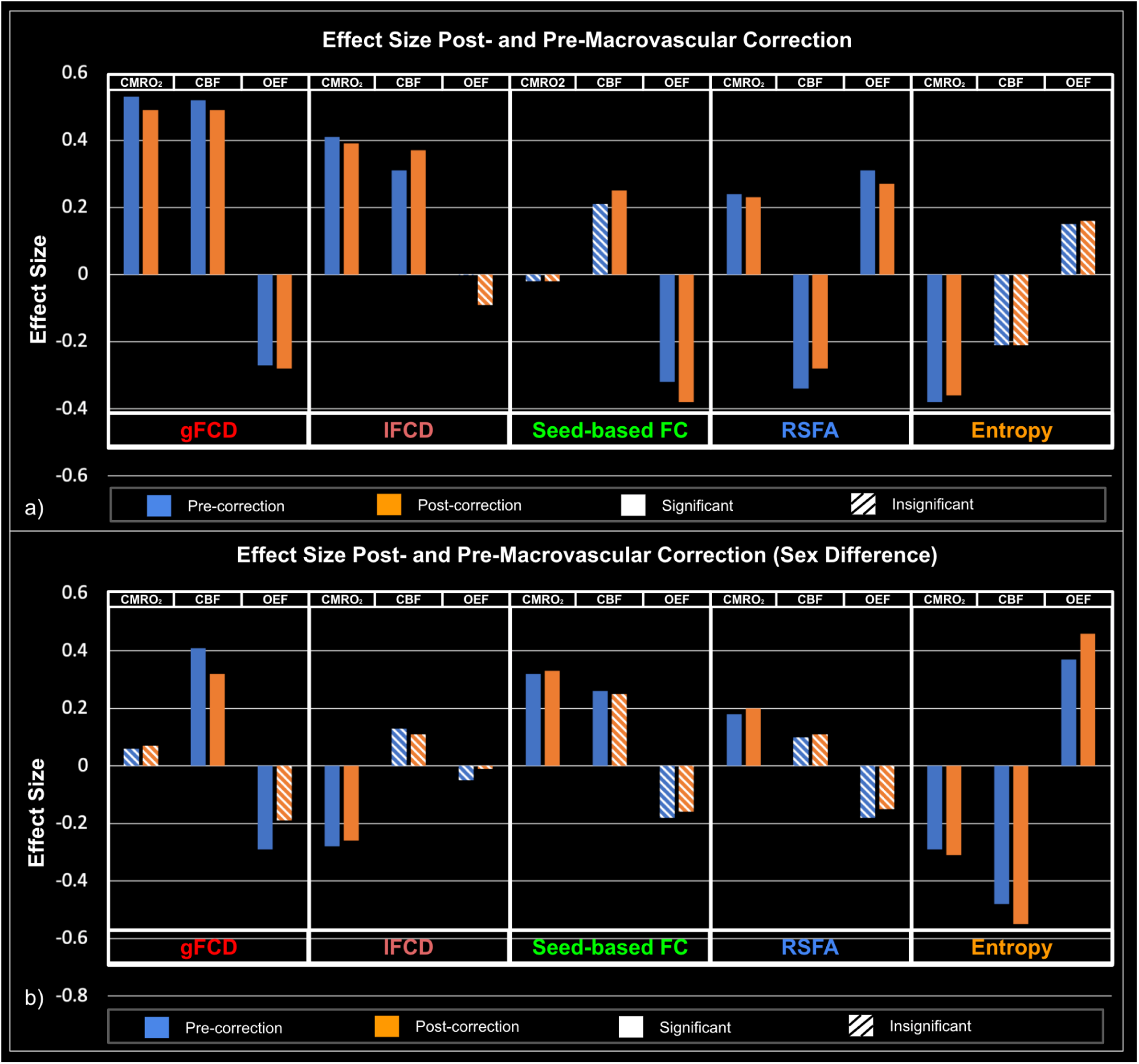
Associations between rs-fMRI metrics and baseline physiological variables (OEF, CBF and CMRO_2_), with and without accounting for macrovascular correction. a) Strength of associations between rs-fMRI metrics and baseline physiological variables; b) comparison of strength of associations between males and females, with a positive effect size indicating stronger associations in males. Blue: pre-correction; orange: post-correction; striped bar: significant association; dashed bar: insignificant association.

OEF and CBF (**Fig. 4a**) were found to be significantly associated with both seed-based and seed-independent rs-fMRI metrics. OEF was negatively associated with gFCD and seed-based FC and positively associated with RSFA, whereas CBF was positively associated with gFCD, lFCD, seed-based FC and negatively associated with RSFA.

We found significant sex differences in terms of the associations with CMRO_2_ (**Fig. 4b**), which suggests the associations between CMRO_2_ and both entropy and lFCD are stronger for females, and the associations between CMRO_2_ and both seed-based FC and RSFA are stronger for males. We also found significant sex differences in terms of the associations with CBF, which suggests the association between CBF and entropy is stronger for females and associations between CBF and gFCD are stronger for males. Moreover, the association between OEF and rs-fMRI entropy is significantly stronger for males. The results presented here pertain to macrovascular-corrected metrics. The next sections consolidate the understanding of the effect of the macrovascular correction.

### The effect of macrovascular correction

#### Comparison of rs-fMRI-physiology associations post and pre-macrovascular correction

Figure 4 illustrates how the significance of the LMEs linking rs-fMRI and physiological variables was altered due to macrovascular correction. Specifically, 1) the association between seed-based FC and CBF became significant only after macrovascular correction; 2) the sex effect in the association between OEF and gFCD became insignificant after macrovascular correction; 3) the sex effect in the association between seed-based FC and CBF became insignificant after macrovascular correction.

#### Goodness-of-fit comparison between post- and pre-macrovascular correction

As shown in Figure 5, in all relationships, the goodness of fit to the LME (as measured by the r^2^) increased when the rs-fMRI metric was based on macrovasculature-corrected data. Notably, the lFCD and RSFA fittings showed the greatest improvement (Fig. 5b), while seed-based FC and entropy fitting (only to CMRO_2_) showed the smallest improvement. Overall, the gFCD-physiology associations were associated with the highest r^2^ of fittings, both pre- and post-correction, followed by the RSFA-physiology associations, with the physiological associations with seed-based FC producing the lowest r^2^ (Fig. 5a). Additionally, as physiological variables go, CMRO_2_ presented a greater r^2^ when fit against rs-fMRI metrics, higher than CBF and OEF.

**Figure 5.**
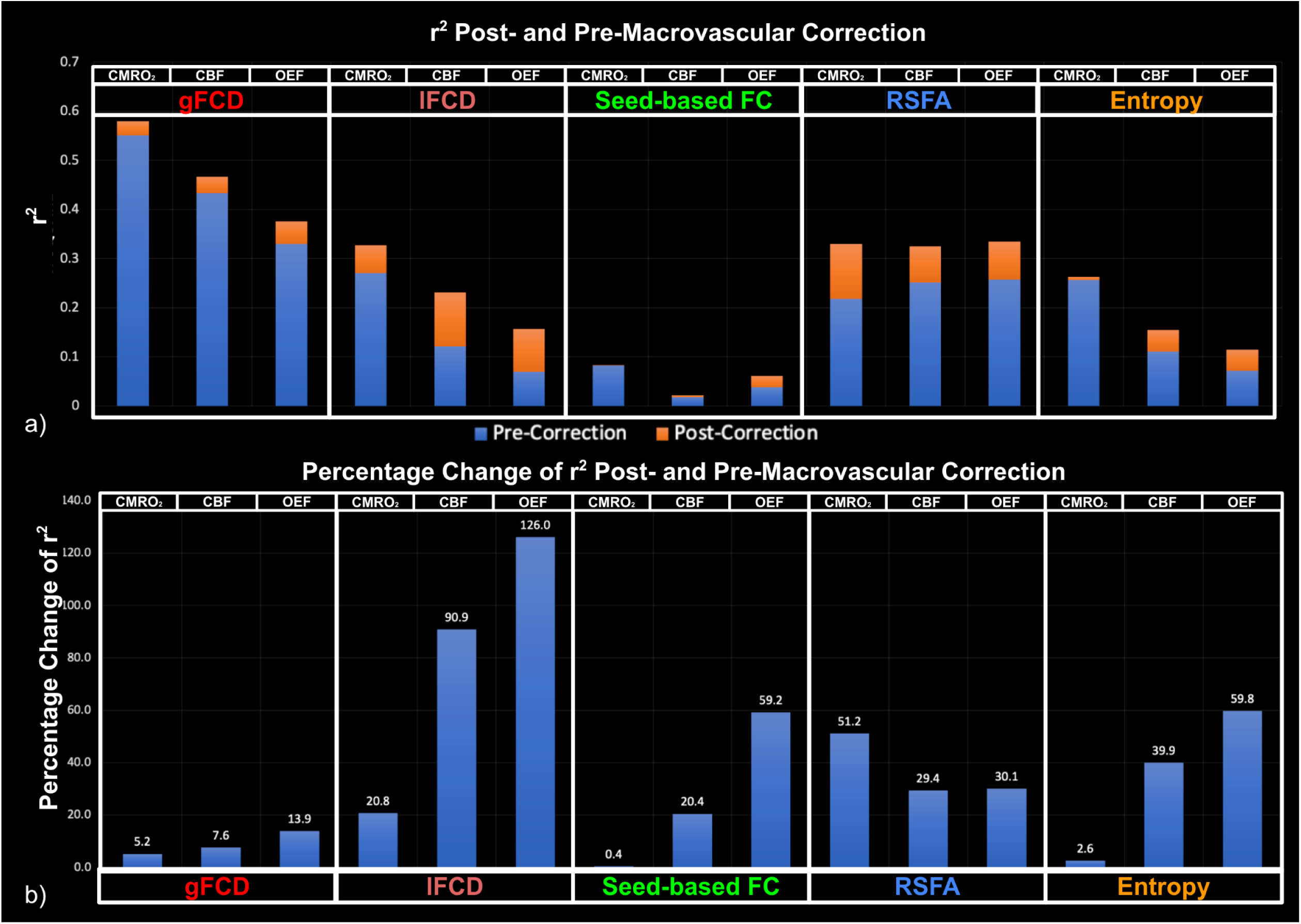
Goodness of fit (r^2^) of LME models, compared between post- and pre-macrovascular correction. a) r^2^ post- and pre-macrovascular correction; b) percentage difference in the r^2^ post- and pre-macrovascular correction.

#### Comparison of rs-fMRI metrics post- and pre-macrovascular correction

To better understand the sources of these differences, Figure 6 further quantifies the effect of macrovascular correction on the values of rs-fMRI metrics. The gFCD difference can be seen globally in this representative dataset, with most voxels displaying a decrease in gFCD post-correction (Fig. 6a); the peak of the decreases overlap with the venous vasculature (refer to Fig. 2, especially the superior sagittal sinus). There was also a post-correction decrease in lFCD in most of the voxels (Fig. 6b), but the spatial extent of the decrease was much smaller than for gFCD. A decrease in RSFA occurred after macrovascular correction, primarily in areas surrounding the macrovasculature (Fig. 6c). rs-fMRI entropy varied in both directions post-correction, with a similar spatial extent as that of RSFA, i.e. limited to the area surrounding the vasculature (Fig. 6d).

**Figure 6.**
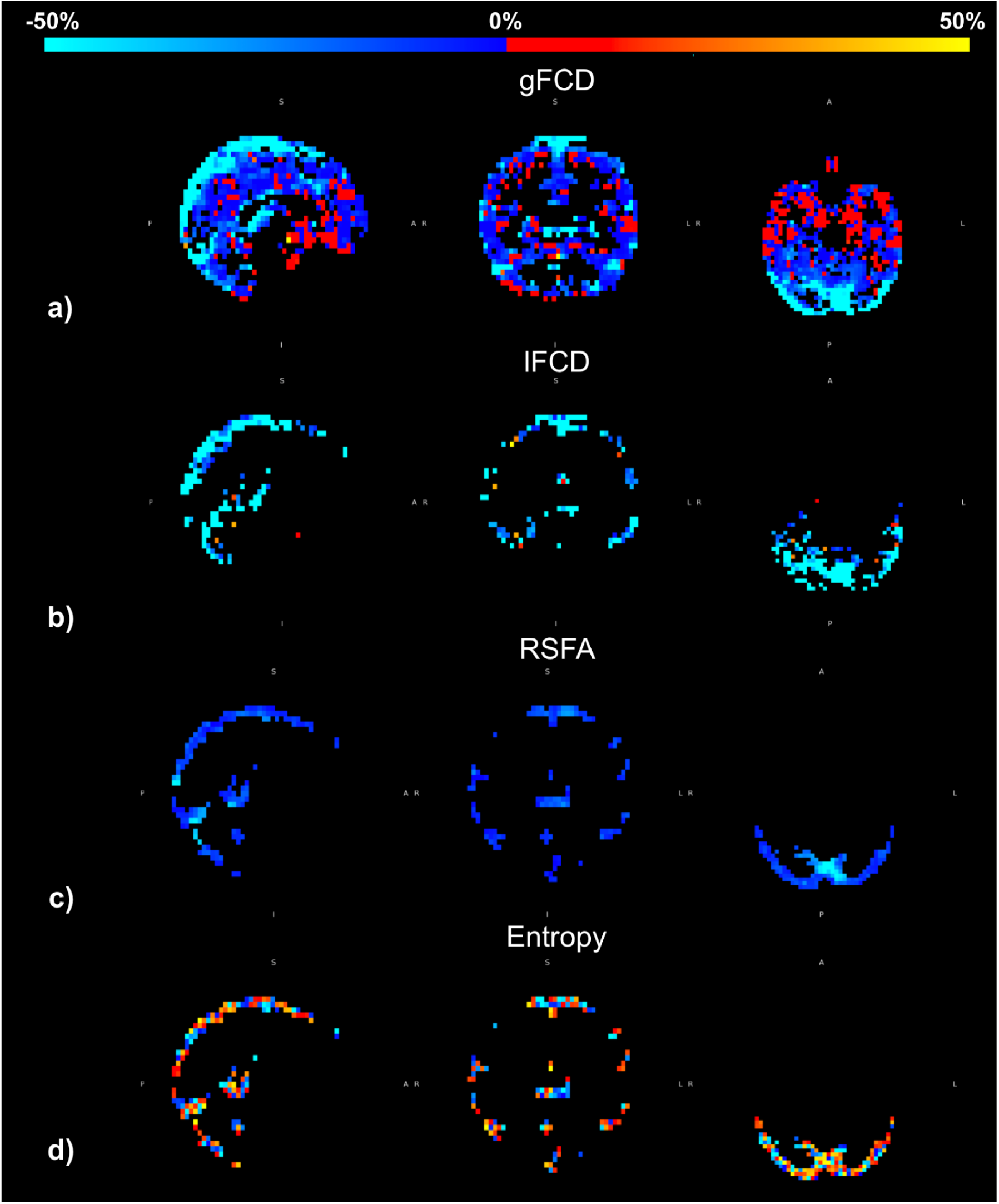
rs-fMRI metrics (network-independent) change after macrovascular correction (Data shown from a representative dataset). a) gFCD; b) lFCD; c) RSFA; d) entropy. The colour bars represent the percent of change (i.e. (post-pre) as a fraction of the post-correction values.

As described earlier, the network ROI definitions depended on whether macrovascular correction was applied to rs-fMRI data. Thus, to help explain any differences in our findings pre- and post-correction, we also examined differences in the network definitions associated with pre- and post-correction fMRI data. As Fig. 7 shows, the spatial extents of the networks extracted pre- and post-macrovascular correction (from a representative participant) showed minimal differences.

**Figure 7.**
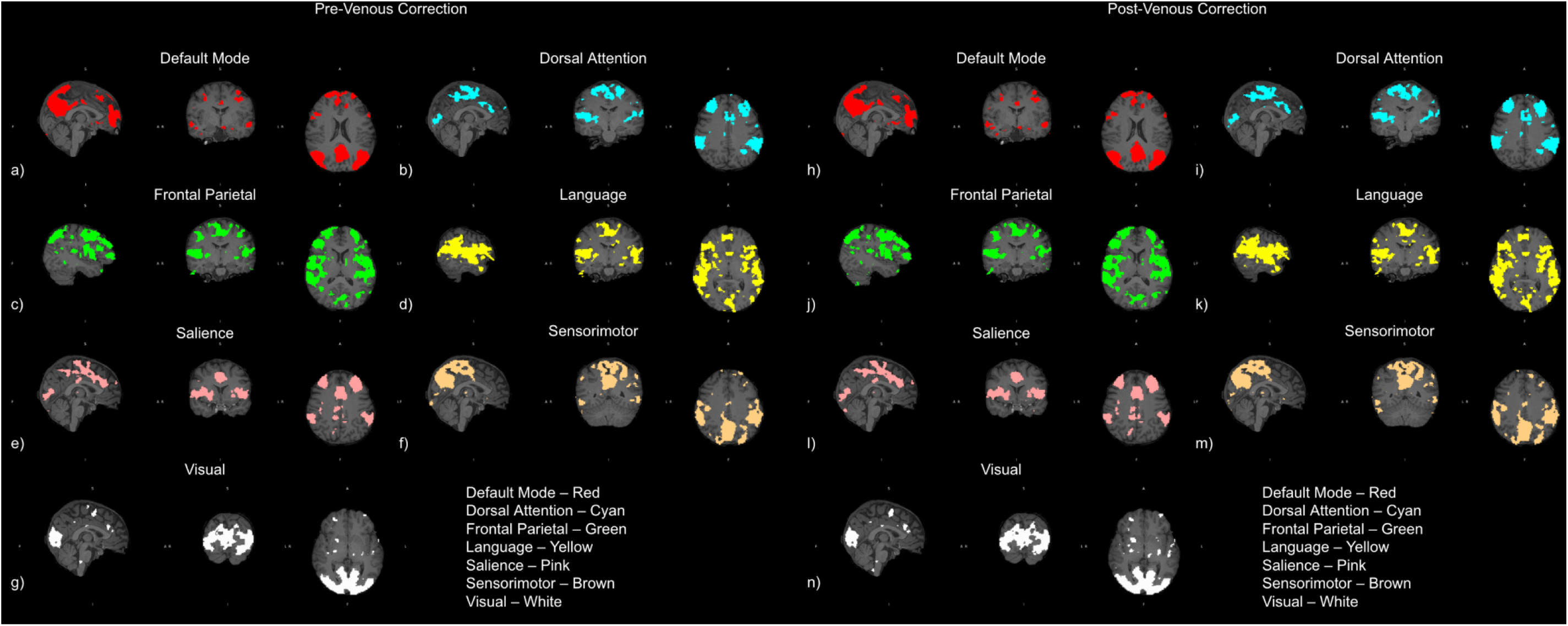
Networks extracted pre- and post-macrovascular correction (Data shown from a representative dataset).

## Discussion

The associations between rs-fMRI metrics and fundamental physiological metrics provide a more solid foundation for understanding rs-fMRI findings. Nevertheless, the presence of the venous bias may obscure the associations. As we demonstrated in our previous research, BOLD signal contributions specific to the macrovascular system (particularly the venous system) can be predicted using biophysical modeling (Zhong, Polimeni, et al., 2024). In this work, we leveraged this biophysical-modeling approach to minimize the macrovascular contributions, and thereafter we found that:

1. The rs-fMRI metrics significantly associated with CMRO_2_ include:

a. gFCD, lFCD and RSFA, which are positively associated with CMRO_2_;
b. rs-fMRI entropy, which is negatively associated with CMRO_2;_
2. The rs-fMRI metrics significantly associated with CBF include:

a. gFCD, lFCD and seed-based FC, which are positively associated with CBF;
b. RSFA, which is negatively associated with CBF;
3. The rs-fMRI metrics significant associated with OEF include:

a. RSFA, which is positively associated with OEF;
b. gFCD and seed-based FC, which are negatively associated with OEF;
4. The use of macrovascular correction substantially increased the goodness-of-fit of all LME models.

### Neuronal activity versus oxidative and glucose metabolism

Previous studies already demonstrated the associations between the rs-fMRI metrics and CMR_glu_ (Shokri-Kojori et al., 2019; Tomasi et al., 2013), but the link between CMRO_glu_ and the BOLD signal is less direct, as similar BOLD responses may be associated with distinct CMR_glu_ responses (Hahn et al., 2024). While oxygen and glucose both serve as fuel for neuronal activity, cerebral glycolytic and oxidative metabolism may not be equally involved. Aerobic metabolism, which produces 38 mol of adenosine triphosphate (ATP) per mol of glucose, is the most efficient way to produce ATP (Krebs & Henseleit, 1932; Morrill, 2021). The entire pathway is composed of two steps: glycolysis in the cytoplasm and Kreb’s cycle in the mitochondria. Glycolysis oxidizes the glucose to pyruvate and produces 2 mol of ATP. In anaerobic conditions, pyruvate is further converted into lactate (Melkonian & Schury, 2024); in aerobic conditions, pyruvate enters the mitochondria and is fully oxidized to carbon dioxide and water through Kerb’s cycle, producing 36 mol ATP. When neuronal activity is in steady state and under normal physiology, it is estimated that 90% of glucose consumption in the brain occurs through oxidative metabolism (molar oxygen–glucose ratio is 5.6) (Gjedde, 2001). However, it is not yet clear whether the coupling between CMR_glu_ and CMRO_2_ would operate in a “steady state” when in resting state.

According to work by Raichle et al. based on positron emission tomography, the DMN exhibits a higher resting-state CMRO_2_ than the rest of the brain (Raichle et al., 2001). However, the same cannot be said for CMR_glu_ (Markello et al., 2022). In fact, resting-state CMRO_2_ and CMR_glu_ maps are distinct from one another (Markello et al., 2022). One of the sources of this discrepancy between observed levels of CMR_glu_ and CMRO_2_ metabolism can potentially be explained by aerobic glycolysis (Vaishnavi et al., 2010), which in this context refers to the use of glucose in excess of that required for oxidative phosphorylation despite the presence of sufficient oxygen to fully metabolize glucose. Specifically, the area of highest aerobic glycolysis in the resting state was found to overlap with the DMN, which is the most CMRO_2_-demanding region of the brain during the resting state as mentioned earlier, suggesting a mismatch between CMRO_2_ and CMR_glu_. It is evident that CMR_glu_ and CBF are linked, but a decrease in CBF does not alter CMR_glu_ (Cholet et al., 1997). To summarize, the relationship between rs-fMRI and CMRO_2_, which we focus on in this study, is likely distinct from that with CMRO_glu_.

### Associating whole-brain rs-fMRI metrics with CMRO_2_ and its correlates

The main findings are based on inter-participant differences, as was the case for previous studies relating rs-fMRI to CMRglu (Shokri-Kojori et al., 2019; Tomasi et al., 2013).

#### Seed-based FC

We found a significant positive relationship between CBF and seed-based FC, consistent with previous research (Chu et al., 2018; Liang et al., 2013). However, the association between FC and CMRO_2_ was not detected, potentially due to the limited range of FC strength (instead of density) across subjects, attributable to the averting of different seed-based FC values (Cole et al., 2010). Moreover, seed-based FC is based on averaged correlation coefficients between seeds within a network, while the seed signal is the averaged signal for a certain brain region (seed region) rather than a voxel-specific signal. Therefore, seed-based approaches may naturally have less local specificity and have a lower association with CMRO_2_, which reflects local-specific neuronal activity.

#### Seed-independent FC: gFCD and lFCD

As FCD is a measure of how many voxels are connected to a given voxel, and thus the degree of coordination between voxels, regions with a high FCD value are considered connectivity hubs (Tomasi & Volkow, 2010, 2011). It is expected that the maintenance of a higher FCD may require more energy (Raichle & Mintun, 2006). This is indeed what we find, in that higher gFCD and lFCD are associated with higher CMRO_2_. This is consistent with the associations between FCD and CMR_glu_. Tomasi and colleagues demonstrated a significant positive correlation between CMRO_glu_ and lFCD and gFCD in the DMN, the DAN, and the cerebellar network (Tomasi et al., 2013) and across grey matter (Bernier et al., 2017; Shokri-Kojori et al., 2019). Our study goes a step further by evaluating the link between gFCD, lFCD and CMRO_2_, while also finding a positive relationship with CBF and a negative one with OEF.

We noticed that the association between CMRO_2_ and FCD (both gFCD and lFCD) is more likely to be driven by CBF, since OEF is negatively associated with FCD. That is, the increased metabolic demands caused by a higher resting-state FCD (both global and local) are overcompensated by an increase in CBF, as is the case in task-related BOLD (Buxton et al., 2004). Though we previously stated that CMRO_2_ and CMR_glu_ might not behave similarly in resting state, they appear to exhibit similar associations with gFCD and lFCD). As such, resting-state cerebral metabolism can be viewed as a form of steady state, as defined previously (in terms of the potential links between CMRO_2_ and CMR_glu_) (Gjedde, 2001).

#### RSFA

RSFA could reflect both strong neuronal (Leopold et al., 2003; Zhong & Chen, 2022) and vascular contributions (Chu et al., 2018; Golestani et al., 2016; Tsvetanov et al., 2015), which complicates the interpretation of RSFA. Previous studies have been inconsistent in terms of whether low-frequency BOLD amplitude is associated with CMR_glu_ (Tomasi et al., 2013) or not (Bernier et al., 2017). We found a significant positive association between RSFA and CMRO_2_ in our study, which mirrors findings linking the RSFA and CMR_glu_ as found by Tomasi and colleagues (Tomasi et al., 2013). Therefore, RSFA does not merely reflect vascular activity, but also the metabolism associated with local neuronal activity. Compared to connectivity metrics, the strength of associations between CMRO_2_ (as well as OEF and CBF) and RSFA is lower than FCD metrics, but is still stronger than seed-based FC.

Aside from metabolism, baseline CBF is also reportedly significantly associated with the RSFA, the direction of which varies according to the ROI selected (Chu et al., 2018). Results in this study indicate that RSFA is negatively associated with CBF but positively associated with OEF. This could be driven by the nature of RSFA, which is baseline-dependent. As illustrated by previous simulations (Chu et al., 2018), a higher baseline CBF as well as a higher resultant blood oxygenation level would both lead to a lower RSFA, irrespective of the underlying neuronal activity. Thus, we do not regard the RSFA as a sufficiently specific metric to assess neuronal activity.

#### Entropy

Resting-state fMRI entropy is a relatively new metric, which measures the temporal complexity of signals and has been found to negatively correlate with cognitive ability in healthy young adults (Del Mauro & Wang, 2024a), as well as with cortical thickness and surface area in some brain regions (Del Mauro & Wang, 2024b). In previous studies, it was hypothesized that higher entropy represents lower functional temporal coherence and a higher cognitive reserve (Del Mauro & Wang, 2024a, 2024b), which in turn can further transform into a lower metabolic budget (the higher metabolic reserve). This is consistent with our results in that participants with higher rs-fMRI entropy have lower CMRO_2_. The association between rs-fMRI entropy and CMRO_2_ illustrates the neural basis of this temporal coherence metric. In addition, it was reported that correlations between CBF and rs-fMRI entropy are limited (Song et al., 2019), in broad agreement with our findings here. It should be noted, however, that even though the CBF fitting turned out to be insignificant, the effect size is not negligible. Future research that includes more participants may be necessary to fully comprehend these findings.

Entropy showed one of the strongest associations with CMRO_2_, second only to gFCD, albeit in the opposite direction. However, the association of entropy and CBF is poor, suggesting that entropy offers a different perspective on brain function than conventional connectivity metrics.

### Macrovascular bias and correction in rs-fMRI

Despite the fact that both large veins and large arteries have a strong effect on the functional connectivity of the brain (Tong, Yao, et al., 2019; Zhong, Tong, et al., 2024), the effect of large veins is much greater than the effect of large arteries (Zhong, Tong, et al., 2024). Moreover, predicting the arterial BOLD signal with a biophysical model is more challenging than predicting the venous BOLD signal (Zhong, Polimeni, et al., 2024). Since our biophysical model is most reliable for predicting venous and perivenous BOLD signals, this study only addresses venous bias correction.

The theory behind the macrovascular bias of rs-fMRI metrics can be found in our previous work (Zhong, Polimeni, et al., 2024; Zhong, Tong, et al., 2024). In short, the presence of macrovasculature directly biases the RSFA, which in turn is associated with non-linear biases in other rs-fMRI metrics (Chu et al., 2018; Zhong, Polimeni, et al., 2024). The resultant connectivity is most likely to be overestimated. We used these theoretical predictions to help further establish the validity of the correction, through three main arguments.

First, we confirmed the effect of the correction against theoretical predictions. The correction altered the gFCD, lFCD, and RSFA, as illustrated in Fig. 6. The RSFA and lFCD, as predicted from theory (Chu et al., 2018; Zhong, Polimeni, et al., 2024), decreased. The gFCD also decreases post-correction for the most part. Notably, all reductions closely follow the venous vasculature. However, the gFCD showed a few regional increases as well, which could be the result of a variety of factors. Although the macrovasculature is shown to be strongly correlated within itself (Zhong, Polimeni, et al., 2024; Zhong, Tong, et al., 2024), it may not always add to the overall FC strength. That is, if the macrovascular signal was to interfere destructively with the neuronal signal, then the overall FC may be reduced due to the macrovascular presence. Another factor contributing to a higher gFCD post-macrovascular correction may be artifactual. As the macrovascular correction is based on a single venous regressor, it is conceivable that venous contributions may be artificially introduced into voxels that have none. Moreover, given that BOLD signals from large arteries tend to be strongly anti-correlated with those from the venous vasculature (Zhong, Polimeni, et al., 2024; Zhong, Tong, et al., 2024), gFCD can also increase post-correction in the vicinity of arteries. The increase in network-wise entropy after macrovascular correction is intriguing, suggesting that non-linear temporal complexity metrics are not fully immune to physiological contributions, as has been suggested previously (Del Mauro & Wang, 2024a). Further research may be needed to determine whether the current view regarding the interpretation of rs-fMRI entropy stems from the contamination of macrovascular bias.

The second argument is whether macrovascular correction improved the neuronal specificity of rs-fMRI measurements. It may be possible to answer the question by testing the goodness-of-fit (r^2^) of the association between rs-fMRI metrics and baseline brain physiology metrics, especially CMRO_2_, as explained earlier. A higher r^2^ after macrovascular correction would suggest that rs-fMRI metrics are closer to their BOLD physiological origins and can better reflect local-specific neural activity (Kim et al., 1999; Raichle et al., 2001) - this is exactly what we observed in all cases (Fig. 5). Therefore, these findings support the use of macrovascular correction in the context of BOLD-based rs-fMRI metrics. They also, establish the metabolic associations with these multiple rs-fMRI metrics in multiple brain regions, substantially extending previous findings based on CMRO_2_ and CMRglu (Raichle et al., 2001; Shokri-Kojori et al., 2019; Tomasi et al., 2013). It appears that lFCD and RSFA have the most significant improvements in terms of associations with each of the rs-fMRI metrics (Fig. 5). Based on our biophysical model, the RSFA can also be strongly affected by this bias (Ogawa et al., 1993; Zhong, Polimeni, et al., 2024). Hence, it is not surprising that associations of RSFA with CMRO_2_ (and its correlates) show strong improvement after macrovascular correction. Moreover, in this regard, the lFCD showed stronger improvements than gFCD, as the definition of the lFCD neighbourhoods favours neighbouring voxels with higher correlation, which fit the behaviour of voxels containing large vessels. In contrast, seed-based FC shows very limited improvement following macrovascular correction (Fig. 5), but as seed-based FC shows an inherently low association with CMRO_2_ and low r^2^ (see Seed-based FC section), it is deemed not the best surrogate for neuronal metabolism based on our findings. The extent to which venous bias affects the measurement of entropy remains unclear, but it is also the case that we do not understand the entropic behaviour of macrovascular BOLD.

The final argument for macrovascular correction is the value added to rs-fMRI analyses as a result of the correction. We provide evidence that certain relationships can only be observed after the correction. For example, the relationship between seed-based FC and CBF is only significant after macrovascular correction has been applied (Fig. 4). It is also important to note that with more participants, the difference between pre- and post-macrovascular correction could be even more evident. Additionally, some associations were eliminated after the macrovascular correction. Some examples include the sex dependence of the gFCD-OEF association and of the seed-based FC-CBF association (Fig. 4). The correction did not strengthen any sex dependence in the rs-fMRI-physiology associations. As we cannot justify the association between rs-fMRI and physiology being sex-dependent, these findings may be a sign that some of the associations that we previously found may have been caused by venous bias rather than neuronal activity.

### Recommendation

Our current study goes beyond our previous studies (Zhong, Polimeni, et al., 2024) by applying the biophysical modelling-based macrovascular correction and validating its effectiveness. We would like to reiterate the recommendations that we made in our previous article that TOF images should be collected along with the BOLD measurements (Zhong, Tong, et al., 2024) and that a macro-VAN model can be used to predict venous behaviour with great accuracy (Zhong, Tong, et al., 2024). Moreover, we show that it is feasible to use the BOLD signal from the voxel with maximum fBV as a surrogate for Yv variation in the simulation of the biophysical model. Furthermore, venous signal contributions could be corrected through regression with a lag shift in the simulated venous signal, as we outlined here. More sophisticated correction approaches (for example, principal component analysis (PCA) and independent component analysis (ICA)) will be examined in future research. Lastly, based on these findings, we recommend that macrovascular corrections be applied to all rs-fMRI experiments focusing on localized neuronal interpretations. More broadly, our methods can be applied to all fMRI studies. However, greater caution needs to be exercised for experiments focusing on more global BOLD effects (such as sleep studies and autonomic nervous system studies), as the interpretation of the macrovascular signals may be less clear in such cases.

### Limitations

Limitations directly related to TOF imaging acquisition and marco-VAN model simulation have already been addressed in our previous paper (Zhong, Polimeni, et al., 2024; Zhong, Tong, et al., 2024) and will not be repeated here. We want to emphasize that we did not include arterial bias in the correction. Moreover, this study focuses on the feasibility and validation of our macrovascular correction approach, and thus does not include a comprehensive comparison with previous macrovascular correction approaches. While we advocate for macrovascular correction on the merits of a stronger resultant association with CMRO_2_, macrovascular contributions to CMRO_2_ cannot be completely eliminated. MRI-based CMRO_2_ measurements are also sensitive to macrovascular contributions, considering they are also based on R_2_’ contrast. Although the OEF estimates are based on bias-field corrected R_2_’ estimates, the effect of venous bias could remain. Thus, while using CMRO_2_ and OEF to test the effect of rs-fMRI macrovascular correction is a major step forward in establishing the neuronal implications of rs-fMRI, non-MRI-based validation (for example, using EEG) would also need to be considered.

## Conclusion

Clinical applications of rs-fMRI are challenging as there is no clear understanding of the physiological mechanisms that underlie rs-fMRI metrics. In order to make the rs-fMRI metrics more interpretable, it would be necessary to understand the relationship between rs-fMRI metrics and CMRO_2_, which is directly related to neuronal activity. This study helps establish these associations across multiple brain regions, as well as demonstrates how our novel macrovascular correction can enhance these associations. As we show the interpretability of the rs-fMRI metrics being further improved following macrovascular correction, we suggest that macrovascular correction enhances local-neuronal specificity of rs-fMRI metrics. Results presented in this article may be used to improve the interpretability of rs-fMRI metrics in the future, as well as to promote its clinical applications.

## Acknowledgement

The authors would like to acknowledge financial support from Canadian Institutes of Health Research and the Canada Research Chairs Program (JJC) and funding support from Ydessa Hendeles Graduate Scholarship (XZZ). We extend our special thanks to Dr. Christine Preibisch and her collaborators for their guidance on R_2_’ estimation and special thanks to Dr. Danny J. J. Wang and his collaborators for providing DE-pCASL sequence.

