## Supplemental Figures for "The Link to Oxidative Metabolism Varies across rs-fMRI Metrics: A Whole-Brain Assessment Using Macrovascular Correction"

### Venous signal simulation and correction

A whole-brain susceptibility map was generated numerically using the Fourier method (Eq. 1 and 2), in which a mask of the vasculature was constructed by upsampling macro-VANs to 0.175 mm isotropic resolution and zero-padding by the size of a full field-of-view on each side to avoid wraparounds resulting from cycle convolution (Cheng et al., 2009; Salomir et al., 2003). To calculate the susceptibility difference between blood and tissue outside macrovasculature, we assumed that the tissue type outside macrovasculature is grey matter (GM).

$$\Delta B_z = FT^{-1}[(\frac{1}{3} - \frac{k_z^2}{k^2})FT(\chi)] \quad (1)$$

$$\chi = \Delta\chi \cdot Hct \cdot (1 - Y) \quad (2)$$

FT denotes the Fourier transform, $\chi$ the local susceptibility, $k_z$ the distance in the k space along the z-axis and k the distance in k-space ($k^2 = k_x^2 + k_y^2 + k_z^2$). $R_2'$ is then calculated through the magnitude of the complex-valued mean magnetization of the dephasing spins resulting from the $B_0$ offset.

The mean BOLD signal was calculated as defined by Eq. 3 and 4.

$$S_{T2'} = |\mu(exp(i\gamma\Delta B_z TE))| \quad (3)$$

$$S = sin(\alpha)(1 - exp(-TR/T1))/(1 - cos(\alpha)exp(-TR/T1)\, exp(-TE/T2))S_{T2'} \quad (4)$$

where $\gamma$ is the gyromagnetic ratio, and the operator $|\mu(\cdot)|$ represents the magnitude of the mean of a complex number. A summary of the values and definitions used for simulation parameters is provided in **Table S1**.

The simulations were conducted using our servers equipped with 14 cores of Intel Xeon X5687 CPU (at 3.6 GHz) (Intel Corporation, Santa Clara, CA, United States) and 180 GB of memory running Red Hat Enterprise Linux Server 7.7 (Red Hat Inc., Raleigh, NC, United States). A customized simulation script was written in MatLab 2019b (MathWorks Inc., Natick, MA, United States.).

**Table S1. Simulation parameters and values.** For all three simulation models, these values were set to default unless otherwise stated.

| Parameter | Definition | Simulated Value | Source |
|---|---|---|---|
| $\Delta\chi$ | Susceptibility of blood with | $4 \times \pi \times 0.27 \times 10^{-6}$ | (Spees et al., 2001) |

| | | | |
|---|---|---|---|
| | fully deoxygenated blood | | |
| Hct | Hematocrit | 0.4 | Men: 40-54%; Women: 36-48% (Billett, 1990) |
| Voxel size | N/A | 3.5 mm isotropic | According to in-vivo rs-fMRI acquisition protocol |
| TR | Repetition time | 4.5 s | |
| TE | Echo time | 30 ms | |
| α | Flip angle | 90 deg | |
| $B_0$ | Main magnetic field | 3T | |
| $Y_v$ | Venous oxygenation level | 0.6 | (Fan et al., 2014) |
| $Y_{tissue}$ | Tissue oxygenation level | 0.85 | (Gagnon et al., 2015) |
| $T1_{blood}$ | T1 of blood | 1649 ms | (Zhang et al., 2013) |
| $T1_{tissue}$ | T1 of tissue | 1465 ms | (Shin et al., 2009) |
